# Emergence of microbial diversity due to cross-feeding interactions in a spatial model of gut microbial metabolism

**DOI:** 10.1101/059956

**Authors:** Milan JA Van Hoek, Roeland MH Merks

## Abstract

**Background:** The human gut contains approximately 10^14^ bacteria, belonging to hundreds of different species. Together, these microbial species form a complex food web that can break down food sources that our own digestive enzymes cannot handle, including complex polysaccharides, producing short chain fatty acids and additional metabolites, *e.g.*, vitamin K. The diversity of microbial diversity is important for colonic health: Changes in the composition of the microbiota have been associated with inflammatory bowel disease, diabetes, obestity and Crohn’s disease, and make the microbiota more vulnerable to infestation by harmful species, *e.g.*, Clostridium difficile. To get a grip on the controlling factors of microbial diversity in the gut, we here propose a multi-scale, spatiotemporal dynamic flux-balance analysis model to study the emergence of metabolic diversity in a spatial gut-like, tubular environment. The model features genome-scale metabolic models of microbial populations, resource sharing via extracellular metabolites, and spatial population dynamics and evolution.

**Results:** In this model, cross-feeding interactions emerge readily, despite the species’ ability to metabolize sugars autonomously. Interestingly, the community requires cross-feeding for producing a realistic set of short-chain fatty acids from an input of glucose, If we let the composition of the microbial subpopulations change during invasion of adjacent space, a complex and stratifed microbiota evolves, with subspecies specializing on cross-feeding interactions via a mechanism of compensated trait loss. The microbial diversity and stratification collapse if the flux through the gut is enhanced to mimic diarrhea.

**Conclusions:** In conclusion, this *in silico* model is a helpful tool in systems biology to predict and explain the controlling factors of microbial diversity in the gut. It can be extended to include, *e.g.*, complex food source, and host-microbiota interactions via the gut wall.

## Background

The human colon is a dense and diverse microbial habitat, that contains hundreds of microbial species [1]. These species together form a food web that breaks down complex polysaccharides into monosaccharides and these monosaccharides into short chain fatty acids (SCFAs). The diversity and composition of the intestinal microbiota is correlated with a human health and disease [2]. The composition of themicrobiota is correlated with obesity and inflammatory bowel disease [3] microbial diversity is lower in obese then in lean individuals [4], and patients with diarrhea-predominant irritable bowel syndrome show large temporal shifts in the composition of the micobiota [5]. Metabolic interactions are probably the most important source of bacterial diversity in the colon [6]. The main food sources entering the colon are non-degraded polysaccharides, including resistant starch and cellulose, oligosaccharides, proteins and simple sugars [7]. In addition to these exogenous sources of sugar, the colon wall secretes mucins, which are an important food source [7] for the microbiota.

In paper we ask what mechanisms are responsible for the diversity of the gut microbiota. The structured environments and and diversity of undigested food sources (*e.g.,* complex polysaccharides, *e.g.*, found in food fibers) found in the gut have been shown to sustain diverse microbial communities [2, 8], but interestingly diverse ecosystems can already arise in an unstructured environment with only one primary resource [9, 10, 11, 12, 13]. In long-term evolution experiments of *E. coli* in glucose-limited continuous culture, such emergence could be attributed to crossfeeding interactions, in which one population degrades the primary food source into a secondary food source, thus creating a niche for the second population [12]. Mathematical modeling can help understand under what conditions cross-feeding in microbial communities can emerge and drive diversification of the population. In their isologous diversification model, Kaneko and Yomo [14, 15] studied sets of identical, chaotically oscillating metabolic networks that exchange metabolites via a common, shared medium. Although small populations of oscillators will easily synchronize with one another, larger populations will break up in specialized, synchronized sub-populations. A number of authors asked when such specialization and cross-feeding becomes evolutionary beneficial. Cross-feeding can evolve if there exists a trade-off between uptake efficiency of the primary and secondary food source [16], if a trade-off between growth rate and yield [17], or, in absence of metabolic trade-offs, if the enzymatic machinery required to metabolize all available nutrients is so complex that distribution of enzymes across species in conjunction with cross-feeding becomes the more probable, ‘easier’ evolutionary solution [18].

These initial theoretical and computational models included simplified or conceptual models of metabolism. More recently, it has become feasible to construct dynamical models of microbial communities based on genome-scale metabolic network models (reviewed in Ref. [19]). In these models, multiple species of bacteria interact with one another by modifying a common pool of metabolites. These models are based on (static optimization-based) dynamic flux-balance analysis (dFBA) [20], coupling the optimization-based flux-balance analysis (FBA) approach to modeling intracellular metabolism, with an ordinary-differential equation model (ODE) for the metabolite concentrations in the substrate. These community models more closely approximate microbial metabolism than the initial, more abstract models, such that the results can be compared directly to experimental observations. For example, Tzamali and coworkers [21] used multispecies dFBA to compare the performance of metabolic mutants of E. coli in batch monoculture versus its performance in co-culture with an alternative mutant. Their model predicted co-cultures that were more efficient than their constituent species. Louca and Doebeli [22] proposed methodology to calibrate the bacterial models in such dynamic multispecies
FBA approaches to data from experimental monocultures. By coupling these calibrated dynamics models of isolated strains of *E. coli*, the framework could reproduce experimentally observed succession of an ancestral monoculture of *E. coli* by a cross-feeding pair of specialists. Because these models assume direct metabolic coupling of all species in the model via the culture medium, the model best applies to well-mixed batch culture systems or chemostats. The more recent coupled dynamic multi-species dFBA and mass transfer models [23, 24, 25, 19], or briefly, spatial dFBA (sdFBA) models are more suitable for modeling the gut microbiota. These spatial extensions of the multispecies dFBA approach couple multiple dFBA models to one another via spatial mass transport models (based on numerical solutions of partial-differential equations), such that bacteria can exchange metabolites with their direct neighbors.

In order to explore whether and under which circumstances a diverse microbial community can arise from a single food source in the gut, here we extended the sdFBA approac to develop a multiscale model of collective, colonic carbohydrate metabolism and bacterial population dynamics and evolution in a gut-like geometry. To this end, we combined spatial models of population dynamics with genome-scale metabolic models for individual bacterial species and a spatial mass transport model. In addition to the sdFBA approaches, we extended the model with an “evolutionary” component, in order to allow for unsupervised diversification of the microbial communities. We inoculate the metabolic system with a meta-population of bacteria containing a set of available metabolic pathways. When, depending on the local availability of nutrients, the bacterial population expands into its local neighborhood the metapopulation gains or looses metabolic pathways at random. We find that spatially structured, microbial diversity emerges spontaneously emerge in our model on a single resource. This diversity depends on interspecies cross-feeding interactions.

## Results

A full multiscale model of the metabolism of the human gut would need to include around 10^14^ individual bacteria belonging to hundreds of bacterial species, for which in most cases genome-scale metabolic models are unavailable. We thus necessarily took a more coarse-grained approach, while retaining biological realism by using realistic genome-scale metabolic network models. We first asked to what extent cross-feeding can emerge in large communities of interacting and diversified bacteria, such as those found in the colon, using a dynamic-species metabolic modeling (DMMM) approach [21, 26, 19], which is an extension of the dynamic flux-balance analysis (dFBA) method [27, 20]. We next asked to what extent spatially diversified microbial communities can emerge in a tube-like environment, if the microbial communities are allowed to specialize to the local availability of metabolites.

### Construction of metabolic model representing a subset of the gut microbiota

We first constructed a hypothetical, but biologically-realistic “supra-organism” model [3, 28], called “metabacterium” here, that represents a sample of the gut microbial community in a single metabolic network model. For this preliminary, explorative study we extended a genome-scale metabolic model of *Lactobacillus plantarum* [29], a resident of the colon and a probiotic strain, with four key metabolic pathways of the intestinal microbial community: (1) propionate fermentation, (2) butyrate fermentation, (3) the acrylate pathway and (4) the Wood-Ljungdahl pathway. The model of *L. plantarum* is suitable for FBA [29] and for FBAwMC [30]; in future versions of our framework this network can be replaced by metabolic network models derived metagenomic data [3] as they become available. The current, simplified network contains 674 reactions (Supplementary File 1), and compares well with consensus metabolic networks of carbohydrate fermentation in the colon [31, 32]. For a schematic overview of the key pathways including in the metabolic network, see Figure 1A.

**Figure 1.**
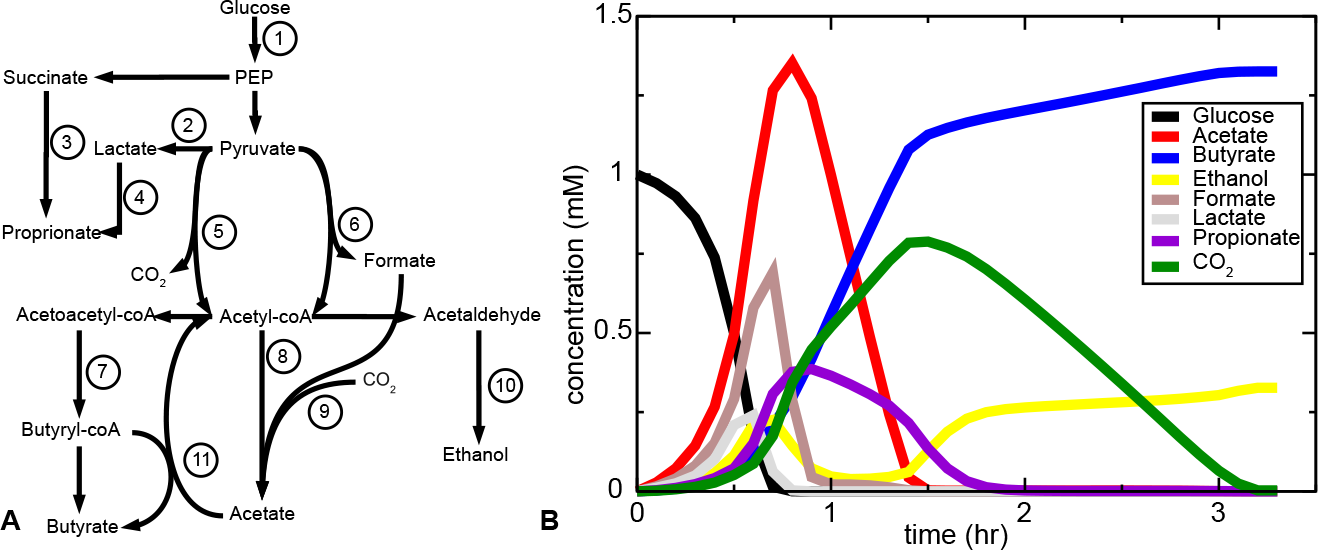
Box initially at rest on sled sliding across ice.A. Simplified scheme of central carbon metabolism of the genome-scale metabolic model. 1) Glycolysis. 2) lactate fermentation. 3) Propionate fermentation. 4) Acrylate pathway. 5) Pyruvate dehydrogenase. 6) Pyruvate formate-lyase. 7) Butyrate fermentation. 8) Acetate fermentation. 9) Acetogenesis via Wood-Ljungdahl pathway. 10) Ethanol fermentation. 11) butyryl-CoA:acetate-CoA transferase. B. Metabolite dynamics over time. At time 0 only glucose is available.

The uptake and excretion rates of genome-scale metabolic networks can be calculated using constraint-based modeling. To represent diauxic growth, *i.e.*, by-product secretion as a function of extracellular metabolite concentrations, we used an extension of FBA called Flux Balance Analysis with Molecular Crowding (FBAwMC) [33]. As an additional, physiologically-plausible constraint FBAwMC assumes that only a finite number of metabolic enzymes fits into a cell, with each enzyme having a maximum metabolic turnover, V_max_. For each reaction, FBAwMC requires a *crowding coefficient*, defined as the enzymatic volume needed to reach unit flux through that reaction. Each reaction is assigned a “crowding coefficient”, a measure of the protein cost of a reaction: Enzymes with low crowding coefficients have small molecular volume or catalyse fast reactions. Given a set of maximum input fluxes, FBAwMC predicts the optimal uptake and excretion fluxes as a function of the extracellular metabolite concentrations.

FBAwMC predicts diauxic growth and the associated by-product secretion in micro-organisms including *E. coli*, *Saccharomyces cerevisiae* [34], and *L.plantarum* [30]. As FBAwMC optimizes growth *rate*, not growth yield as in standard FBA, it predicts a switch to glycolytic metabolism at high glucose concentrations at which faster metabolism is obtained with suboptimal yield. Its accurate prediction of diauxic growth together with by-product secretion as a function of extracellular metabolite concentrations make FBAwMC a suitable method for a microbial community model.

### Metabolic diversity causes cross-feeding in a well-mixed system

To study the extent of cross-feeding emerging already from a non-evolving metabolic community of “metabacteria”, we first set up a simulation of 1000 interacting metapopulations, where each subpopulation was initiated with a set of crowding coefficients selected at random from an experimentally determined distribution of crowding coefficients of *Escherichia coli* [34, 30], for lack of similar data sets for *L. plantarum*. The simulation was initiated with pure glucose. We then performed FBAwMC on all 1000 metapopulations optimizing for ATP production rate. This yielded 1000 sets of metabolic input and output fluxes, F*i*, and growth rates, *μ*_*i*_ for all 1000 metapopulations. These were used to update the extracellular concentrations, 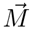 and metapopulation sizes, *X_i_*, by performing a single finite-difference step
of [21, 26]

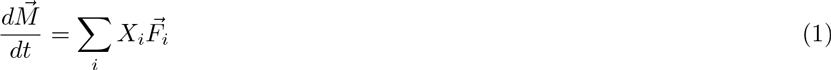

and

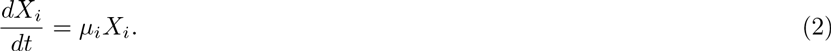

with a timestep Δ*t* = 0.1 h. After updating the environment in this way, we performed a next time simulation step.

Figure 1B shows how, in the simulation, the metabacteria modified the environment over time. The secondary metabolites that were produced mostly are acetate, butyrate, carbon dioxide, ethanol, formate, lactate and propionate. This compares well with the metabolites that are actually found in the colon [35] or in an *in vitro* model of the colon [36]. To test to what extent this result depends on the ability of the individual FBAwMC models to represent metabolic switching and overflow metabolism [33, 30], we also simulated the model using standard flux-balance analysis [37]. In this case, all glucose was converted into ethanol, whereas lactate and propionate did not appear in the simulation (Additional Figure 1). We also studied whether any of the single-species simulations could also provide so many metabolites. Out of 100 single-species simulations none produced as many or more excreted metabolite species than the interacting set of species.

#### Quantification of cross-feeding

Most of the metabolites were only transiently present in the medium, 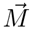, suggesting that the metabolites were re-absorbed and processed further by the bacteria. To quantify the amount of such cross-feeding in the simulations, we defined a crossfeeding factor, *C*(*i*), with *i* a species identifier. Let

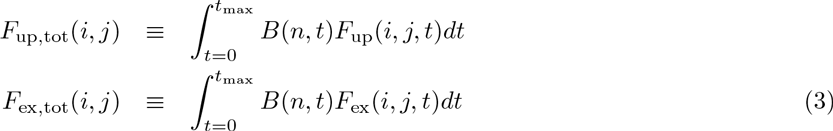

be the total amount of metabolite *j* that species *i* consumes and excretes during the simulation. *B*(*i*,*t*) here equals the biomass of species *i* at time *t*. The amount of carbon species *i* gets via cross-feeding then equals,

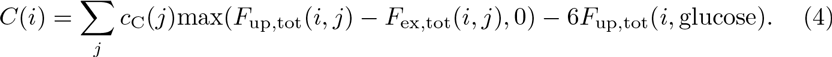

Here, *c*_C_(*j*) is the molar amount of carbon atoms per mol metabolite *j* (*e.g.*, *c*_C_(glucose) = 6). If species i during the fermentation consumes more of metabolite *j* than it has produced, species *i* has cross-fed on metabolite *j*. We subtract the amount of glucose from the sum, because glucose is the primary food source that is present at the start of the simulation. Now we can calculate the total amount of carbon the population acquires via cross-feeding, relative to the total amount of carbon taken up by the population

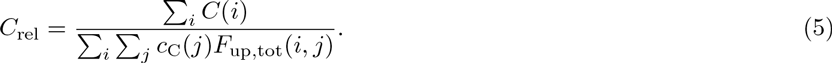

If *C*_rel_ = 0, there is no cross-feeding. In that case, every species only consumes glucose as carbon source or only consumes as much carbon from other metabolites as it has secreted itself. Conversely, if *C*_rel_ = 1 all carbon the species has consumed during the simulation is from non-glucose carbon sources the species has excreted itself. For the whole simulation *C*_rel_ = 0.39 ± 0.02, indicating that 39% of all carbon consumed by the bacteria comes from cross-feeding. Cross-feeding was largest on lactate, CO_2_, acetate, ethanol, formate and propionate. In the original *L. plantarum* model we also find cross-feeding, but only on lactate and acetaldehyde (Additional Figure 2). Taken together, in agreement with previous computational studies that showed cross-feeding in pairs of interacting *E. coli* [21], these simulations show that cross-feeding interactions occur in coupled dynamic FBAwMC models.

### Spatially explicit, evolutionary model

The well-mixed simulations showed that cross-feeding appears in populations of interacting super-organism metabolic networks. However, this does not necessarily imply microbial diversity, because it is possible that the same metabacterium secretes and reabsorbs the same metabolites into the substrate, in which case there would be no true cross-feeding. Furthermore, the previous section did not make clear whether cross-feeding will be ecologically stable under conditions where subpopulations of the supra-organisms are lost. In a spatially explicit model, cross-feeding possibly arises more easily and is more easy to detect, as different metabolic functions can be performed at different locations [38]. We therefore developed a spatially explicit, multiscale evolutionary model of gut microbial metabolism. We initiate the simulation with a population of metapopulations of bacteria that can perform all metabolic functions, just as in the well-mixed simulation. We then let the systems evolve and study if meta-populations of bacteria with specific metabolic roles evolve.

**Figure 2.**
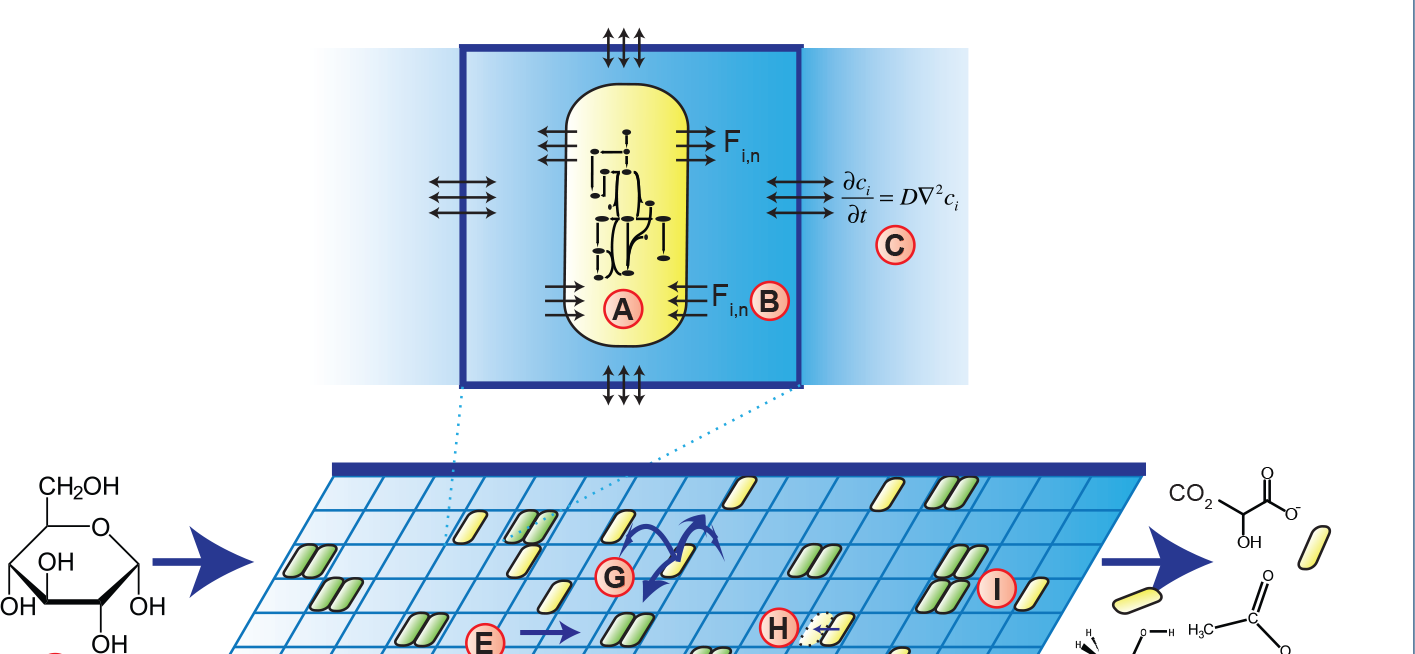
Setup of the simulation model of a metabolizing gut microbial community. The model represents a community of growing subpopulations of genetically identical bacteria. (A) The metabolism of each population is modeled using a genome scale metabolic network model. (B) Based on extracellular metabolite concentrations, the genome scale model predicts the growth rate (*r*) of the subpopulation and the influx and efflux rates of a subset of 115 metabolites. These are used as derivatives for a partial-differential equation model describing the concentrations of extracellular metabolites, 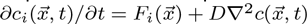, where (C) the metabolites diffuse between adjacent grid sites, 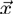. (D) The population is represented on a two-dimensional, tube-like structure, with periodic inputs of glucose. (E) To mimic advection of metabolites through the gut, the concentrations are periodically shifted to the right, until they (F) exit from the end of the tube. (G) The bacterial populations hop at random to adjacent grid sites; to mimic adherence to the gut wall mucus bacterial populations are not advected, unless indicated otherwise. (H) Once the subpopulation has grown to twice its original size, it divides into an empty spot in the same lattice size at which time the metabolic network is mutated. (I) Two subpopulations can live on one grid point; with yellow indicating presence of one subpopulation, and green indicating the presence of two subpopulations. (Structural formulas: Licensed under Public domain via Wikimedia Commons; “Alpha-D-Glucopyranose” by *NEUROtiker*, also licenced under public domain via Wikipedia Commons)

#### Model description

Figure 2 sketches the overall structure of our model. The model approximates the colon as the cross-section of a 150 cm long tube with a diameter of 10 cm. The tube is subdivided into patches of 1 cm^2^, each containing a uniform concentration of metabolites, and potentially a metapopulation of gut bacteria (hereafter called “metabacterium”) (Figure 2A). Each metabacterium represents a small subpopulation (or ‘metapopulation’) of gut bacteria with diverse metabolic functions, and is modeled using a metabolic network model containing the main metabolic reactions found in the gut microbiota, as described above (Figure 1A). Based on the local metabolite concentrations, 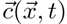 the metabolic model delivers a set of exchange fluxes *F*_i,n_ and a growth rate, 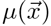, which is assumed to depend on the ATP production rate (Figure 2B; see Methods for detail). The metabolites diffuses to adjacent patches using Fick’s law (Figure 2C), yielding

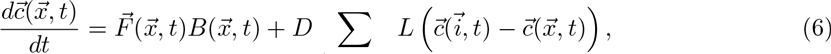

with 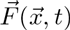 the flux of metabolites between the medium and the metabacterium, the sum running over the four nearest neighbors, 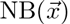, and *L* = 1 cm the interface length between two adjacent patches. The local density of metabacteria, 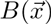 is given by

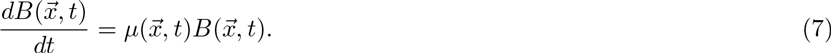

To mimic meals, a pulse of glucose of variable magnitude enters the tube once every eight hours (Figure 2D). The metabolites move through the tube via a simplified model of advection: At regular intervals, all metabolites are shifted one patch (Figure 2E). Metabolites continuously leave the tube at the end through an open boundary condition. To mimic peristaltic movements that locally mix the gut contents together, metabacteria randomly hop to adjacent lattice sites (Figure 2G) and leave the gut only via random hops over the open boundary condition (Figure 2F). In a subset of simulations, accelerated bowel movements are simulated by advecting the metabacteria together with the metabolites. To a first approximation, the boundaries are impermeable to the metabolites, a situation reflecting many *in vitro models* of the gut microbiota (reviewed in Ref.[39]); later versions of the model will consider more complex boundary conditions including absorption of metabolites [40].

When the local biomass in a patch, 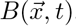, has grown to twice its original value, the metapopulation expands into the second position on the grid point (Figure 2H). To mimic a local carrying capacity, the metapopulation does not spread out or grow any further if both positions in the patch are occupied. In the visualizations of the simulations, full patches are shown in green, singly occupied patches are shown in yellow, and empty patches are shown in black (Figure 2I). During expansion, changes in the relative abundance of species may enhance or reduce the rate of particular reactions, or even delete them from the metapopulation completely. Similarly, metabolic reactions can be reintroduced due to resettling of metabolic species, *e.g.*, from the gut wall mucus [41]. To mimic such changes in species composition of the metapopulation, during each expansion step, we delete enzymes from the metabolic network at random, reactivate enzymes at random, or randomly change crowding coefficients such that the metapopulation can specialize on one particular reaction or become a generalist.

The crowding coefficients, as they appear in the flux-balance analysis with molecular crowding (FBAwMC) method that we used for this model, give the minimum cellular volume filled with enzymes required to generate a unit metabolic flux; they are given by the *V*_max_ of the enzyme and enzyme volume [33]. Equivalently, in our metapopulation model, the crowding coefficient of a reaction is the minimum intracellular volume averaged over all bacteria in the patch that must be filled with enzymes in order to generate a unit flux through the reaction. It depends on the density of the enzyme in the bacteria and also on the corresponding values of *V*_max_. Because the *V*_max_ of a reaction can differ orders of magnitudes between species (see for example the enzyme database BRENDA [42]), the evolutionary dynamics in our model could drive the metabacteria to reduce all crowding coefficients concurrently, producing a highly efficient generalist. To introduce a biologically more realistic trade-off between metabolic rate and cost in terms of volume, we therefore included an experimentally observed trade-off between growth rate and growth yield among micro-organisms [43, 44]: Micro-organisms that grow fast have low growth yield and vice versa. We take this trade-off into account explicitly by assuming a maximal growth rate given the carbon uptake rate of the cells. This trade-off prevents the metabacteria from growing infinitely fast by mutating their crowding coefficients.

**Figure 3.**
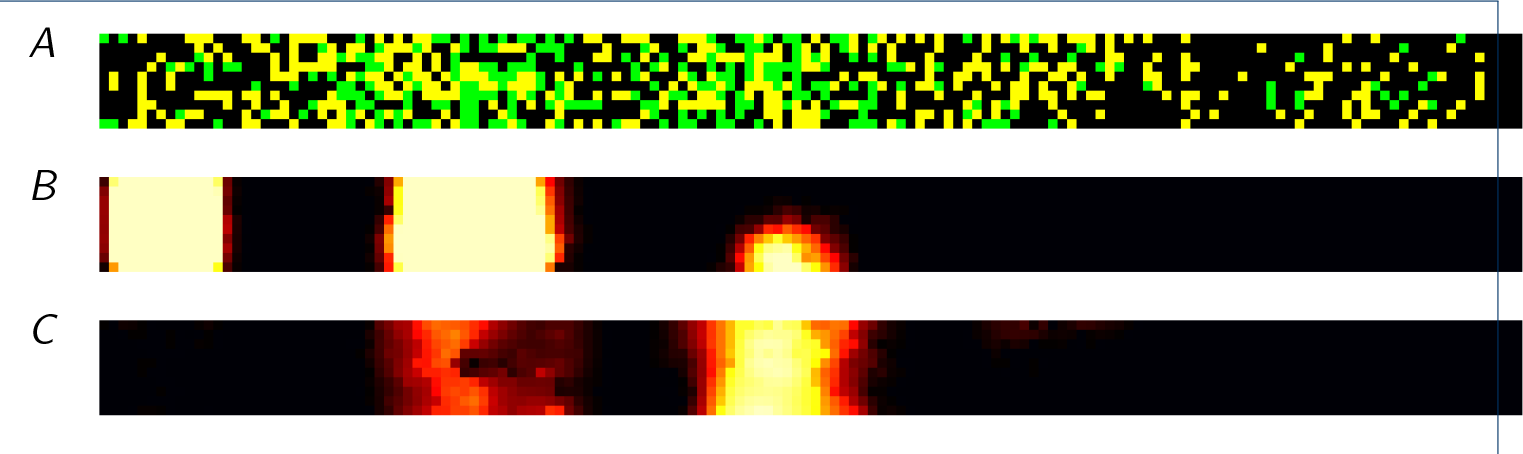
Screenshot of the spatially explicit model. The proximal end of the colon is on the left, the distal end on the right. Thus, food flows from left to right. (*A*) Cells on the grid. At maximum 2 cells can be on the same grid point. Yellow:one cell present, green: 2 cells present. (*B*) Glucose concentration. Black: low concentrations, white: high concentrations. (*C*) Formate concentrations. In total, 115 extracellular metabolites are taken into account in the model.

As an initial condition, we distribute metabacteria over the grid, each containing all available metabolic reactions, *i.e*., each metabacterium initially contains all bacterial “species” that the complete metabacterium represents. To reflect variability in the relative abundances of the bacterial species in each metabacterium the crowding coefficients are drawn at random from an experimental distribution as described above (Figure 2A).

#### Evolution of diversity due to metabolic cross-feeding

To evaluate the behavior of our model, we performed ten independent simulations. These show largely similar phenomenology; therefore we will first describe the progression of one representative simulation in detail, and then discuss differences with the other simulations. Figure 4*A* shows the average number of metabolic reactions present in the metabacteria over time in the simulation. At *t* = 0 all metabacteria still have all 674 reactions, but over time the number of available reactions gradually drops to below 200. This reduction of the number of metabolic genes could indicate a homogenous population specialized, *e.g.*, on fermentation of glucose where in which most of the metabolic network is not used. An alternative explanation is that each of the metapopulation retains a subset of the full network, an indication of cross-feeding. The amount of cross-feeding will likely change over the tube: The metabacteria in the front have direct access to glucose, whereas the metabacteria further down in the tube may rely on the waste-products of those in front. We therefore determined a temporal average of the cross-feeding factors, *C_rel_* (Eq. 5), at each position in the tube over *t* = 3500 to *t* = 4000, a time range at which most genes have been lost. The first observation to note is that in the spatial evolutionary simulations, the average cross-feeding factor *C*_rel_ has a higher value than in the well-mixed simulations. In this particular simulation, the spatial average cross-feeding factor at *t* = 4000 is *C*_rel_ = 0.65 ± 0.09, compared with *C*_rel_ = 0.39 ± 0.02 in the well-mixed case (*n* = 10). The cross-feeding factor for individual cells (*C*(*i*), Eq. 4), showed large population heterogeneity. As Figure 4*B* shows, the cross-feeding factor in the tube front is close to 0, indicating the presence of primary glucose consumers, while cross-feeding slowly increases towards the distal end until it almost reaches 1, indicating complete cross-feeding. Thus in the proximal end the bacteria rely mostly on the primary food source, while near the distal end cells of the tube rely on cross-feeding. This observation is consistent for all simulations (see Additional Figure 3).

**Figure 4.**
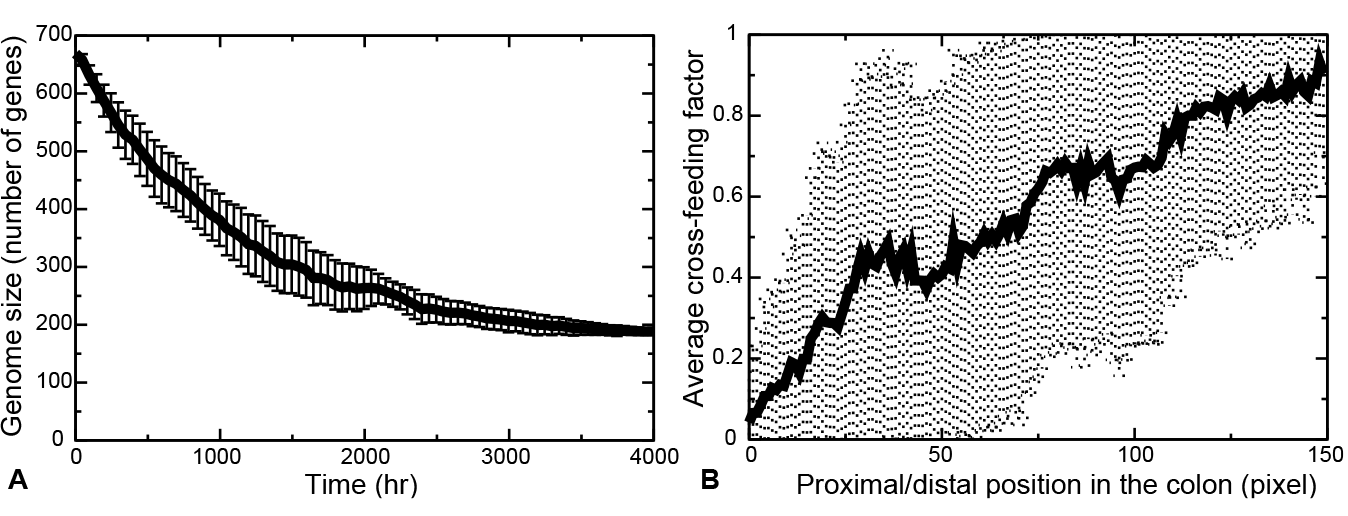
Outcome of the evolutionary simulations. (*A*) population average and standard deviation of the number of enzymatic reactions (“genome size”) over time. (*B*) Population average and standard deviation of the cross-feeding factor *C_n_* as a function of the position in the colon. The averages and standard deviation are over the vertical dimension and are calculated over the final part of the simulation, from 3500 hours until 4000 hours. For the graphs of the other simulations, see Additional Figure 3.

#### Emergence of metabolic stratification

We next investigated the mechanism by which such cross-feeding emerges in the simulation. Additional Figure 4 plots the metabolite concentrations over evolutionary time for the simulation of Figure 4. In this particular simulation, the concentrations of formate and lactate initially rise rapidly, after which they drop gradually. The butyrate concentrations increase over evolutionary time. In all simulations, the metabolite concentrations change gradually, but not necessarily following the same temporal pattern.

Figure 5 shows the spatial distribution of a set of key metabolites averaged over 2000 h to 4000 h of the representative simulation. Interestingly, the flow of metabolites through the colon in interaction with the bacterial population creates a spatially structured, metabolic environment. The proximal end is dominated by the primary carbon source glucose (Figure 5A), with the peak in the average glucose concentration due to the periodic glucose input. Further down in the tube we find fermentation products, including lactate and ethanol, whereas the distal end contains high levels of acetate and CO_2_, showing that the metabacteria convert the glucose into secondary metabolites. Among these secondary metabolites, the levels of acetate (Figure 5B), ethanol (Figure 5E), formate (Figure 5F), lactate (Figure 5G) and propionate (Figure 5H) drop towards the distal end off the tube, so they are further metabolized by the metabacteria. In this particular simulation, butyrate and CO_2_ are not consumed and their concentrations increase monotonically towards the end. The small drop at the very distal end is caused by the metabolite outflow. The profiles of the other simulations were consistent with this representative simulation (Additional Figure 5). In all simulations, the proximal end was dominated by glucose. Further towards the end of the tube, zones of fermentation products developed as in the representative simulation, but the precise location of each product was different and not all products were present. Most notably, in two out of ten simulations, butyrate was absent and in two other simulations proprion-ate was absent. Also, in three out of ten simulations lactate was more confined to the front of the tube (up to around 50 sites) than in the representative simulation.

**Figure 5.**
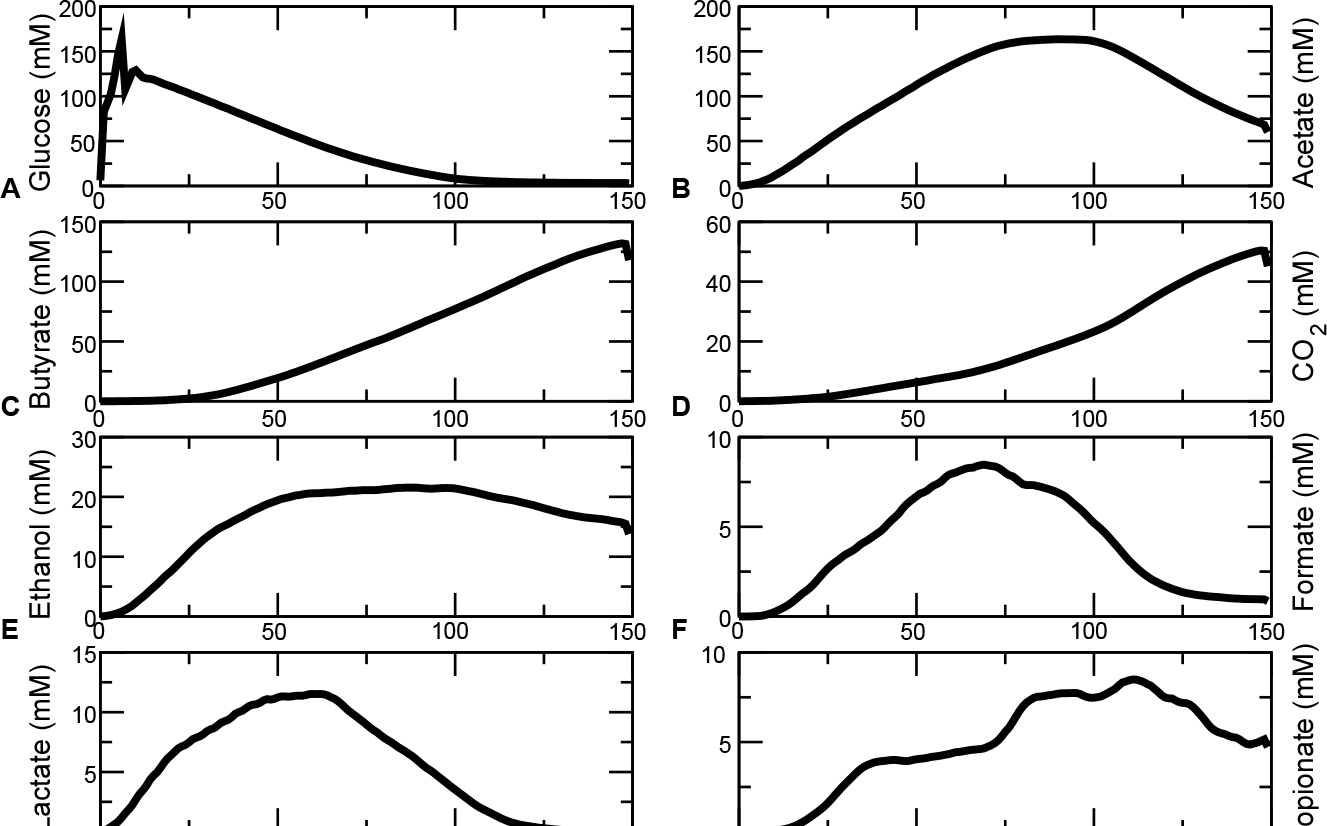
Average metabolite concentrations along the colon. Average are taken over the second half of the simulation (2000hrs-4000hrs). (*A*) Glucose. (*B*) Acetate. (*C*) Butyrate. (*D*) CO_2_. (*E*) Ethanol. (*F*)Formate. (*G*) Lactate. (*H*) Propionate.

#### Metabacteria specialize on local metabolic niches

These results demonstrate that the metabacteria spatially structure their metabolic environment, generating a stratified structure of metabolic “niches” along the tube, each offering a separate set of metabolites. Therefore, we next asked if this environmental structuring gives rise to metapopulations uniquely adopted to the microenvironment. We took computational samples of all metabacteria found in the tube between 3500 h and 4000 h, to average out the variations at the short timescale. We tested the growth rate of these samples (consisting of on average *n* ≈ 1100 metabacteria) in six, homogenous metabolic environments, containing uniform concentrations of pure (1) glucose, (2) acetate, (3) formate, (4) lactate, and (5) propionate, and (6) a mixture of of CO_2_ and H_2_. Figure 6 shows the average and standard deviation of the growth rates of the metabacteria in each of these six environments, as a function of the position from which they were sampled from the tube. Strikingly, the metabacteria near the distal end of the tube have lost their ability to grow on glucose (Figure 6A), indicating that they have specialized on secondary metabolites, including acetate (Figure 6B) and lactate (Figure 6E). Interestingly, in support of the conclusion that the metabacteria specialize on the metabolic niches generated by the population as a whole, the metabacteria sampled from the distal end on average grow faster on acetate and lactate than the metabacteria sampled from the front of the tube. Acetate and lactate are produced in the proximal colon and flow to the distal part of the tube where the metabacteria can metabolize it; in the front of the tube acetate and lactate concentrations are lower, such that neutral drift effects can safely remove the corresponding metabolic pathways from the metabacteria. Remarkably, the metabacteria also grow on CO_2_, because of the presence of hydrogen gas, that allows growth on CO_2_ via the Wood-Ljungdahl pathway [45]. To further characterize the alternative metabolic modes occurring in the model, we clustered the population present at the end of the simulation t = 4000 h with respect to their maximum growth rates in the six environments (Fig. 7). Clearly, different metabolic “species” can be distinguished. One “species” can metabolize glucose, a second “species” can metabolize most secondary metabolites and a third “species” has specialized on acetate. Thus in our simulation model a number of functional classes appear along the tube, each specializing on its own niche in the full metabolic network.

**Figure 6.**
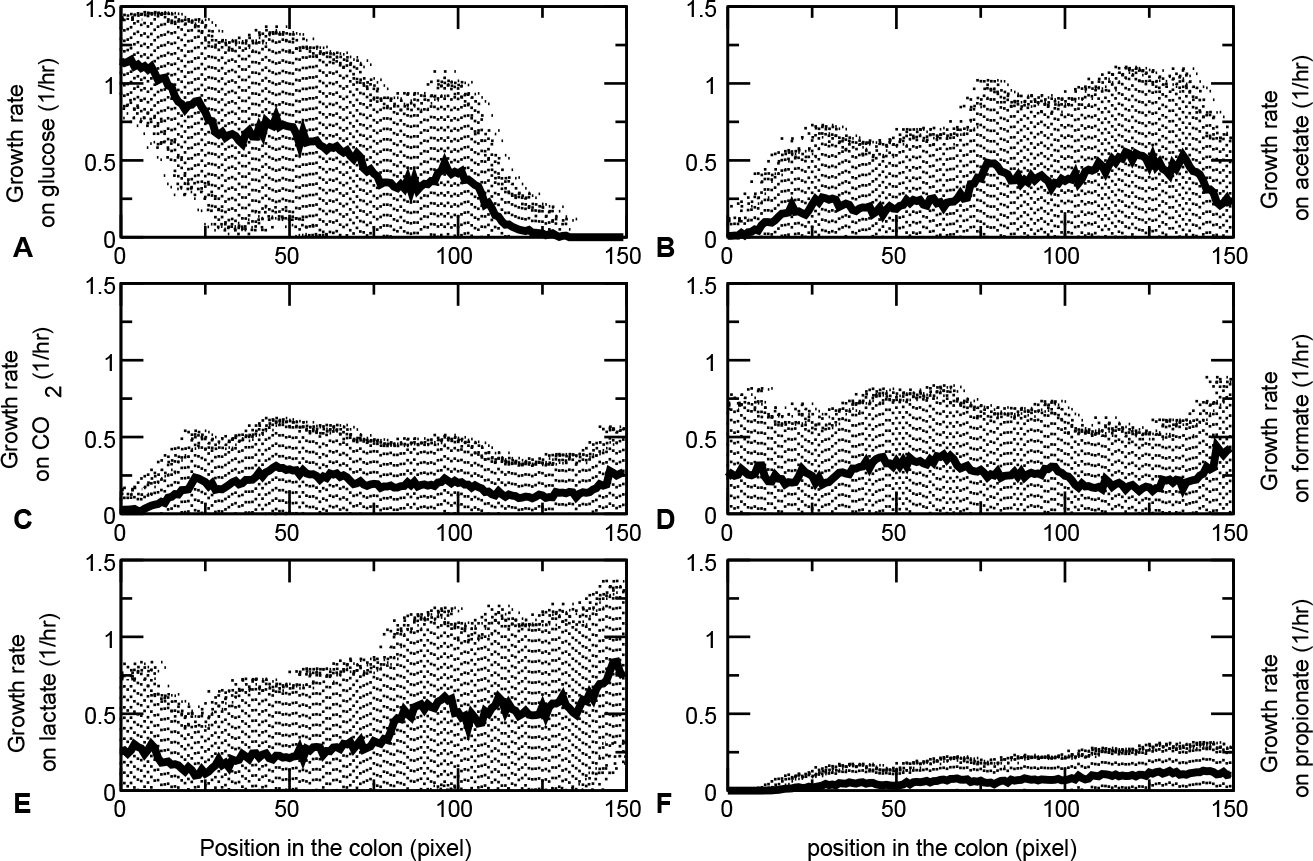
Average growth rates along the colon. Average are taken over the final part of the simulations (3500-4000 hrs) All growth rates ar calculated in the presence of unlimited hydrogen gas, water, sodium, ammonium, phosphate, sulfate and protons. (*A*) Growth rate on glucose. (*B*) Growth rate on acetate. (*C*) Growth rate on CO_2_. (*D*) Growth rate on formate. (*E*) Growth rate on lactate. (*F*)Growth rate on propionate.

#### Increased flux through the tube makes diversity collapse

From the results in the previous section, we conclude that the inherent spatial structuring of the colon results in separate niches. This allows the population to diversify, such that different “species” have different metabolic tasks. A recent population-wide metagenomics study of stool samples from the Flemish and Dutch population [46] showed that, among a range of life-style related factors and medicine use, the diversity of the human gut microbiota correlates strongest with the Bristol stool scale (BSS), a self-assessed indicator of the “softness” of the stool. The analysis showed that for softer stools (higher stool index, indicative of faster transit times [47]), the diversity of the gut microbiota was reduced [48]. To investigate whether transit time could also be correlated with reduced diversity in our model, we studied the effect of increased fluxes through the tube (“diarrhea”), by assuming that the supra-bacteria flow through the tube at the same rate as the metabolites do. Strikingly, the maximal growth rate of the cells has become independent of the position (Fig. 8). Again, we clustered the population present at the end of the simulation with respect to their maximum growth rates in glucose, acetate, H_2_ and CO_2_, formate, lactate and propionate (Fig. 9). In contrast to the simulations without cell flow, the population does practically not diversify. All supra-bacteria can grow on glucose, acetate and H_2_ and CO_2_. Thus, our simulations suggest that increased transit times may contribute to a reduction of microbial diversity, by reducing the spatial heterogeneity in the gut and, consequently, the construction of ecological niches and cross-feeding interactions.

**Figure 7.**
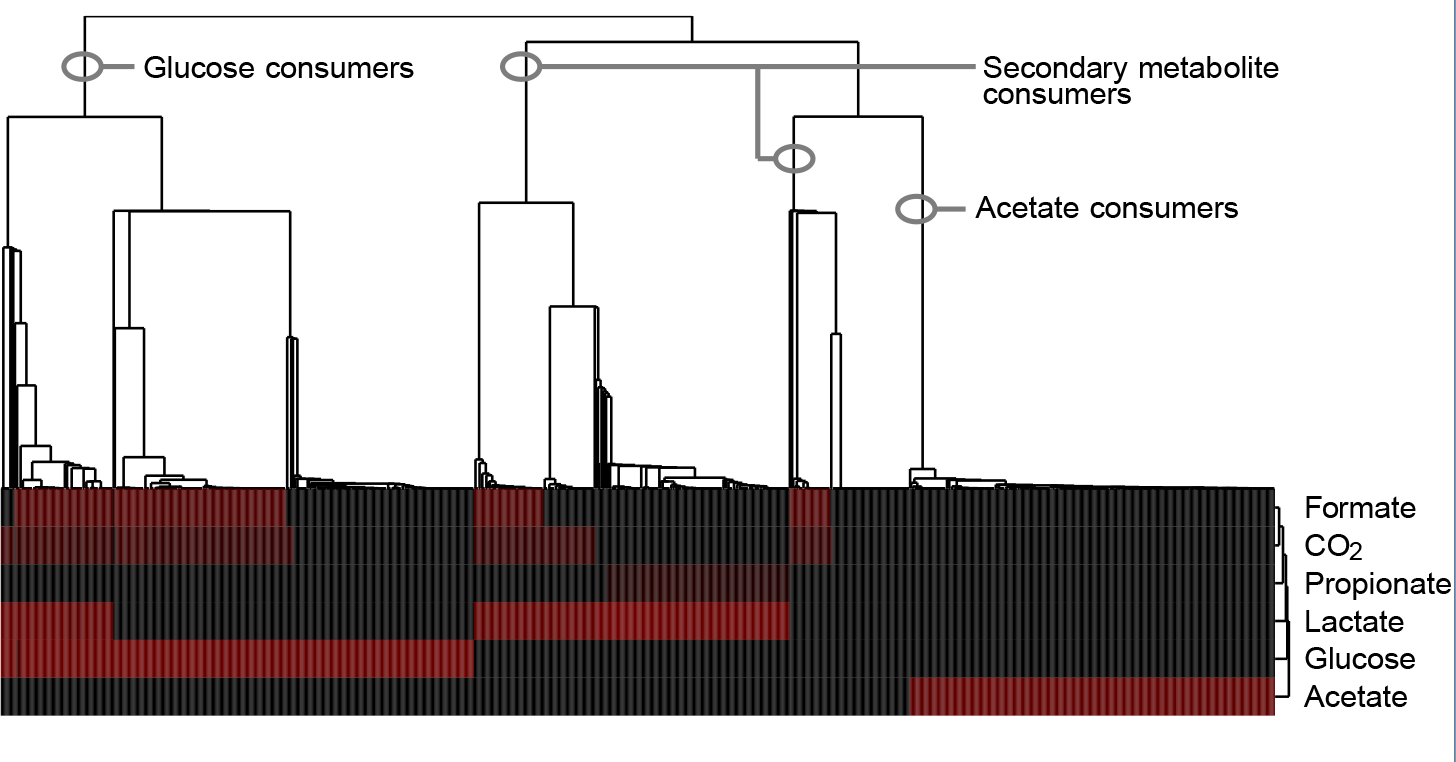
Hierarchical clustering of all cells present at the end of the simulation, with respect to the growth rates on glucose, acetate, CO_2_, formate, lactate and propionate. Black indicates low growth rate, red high growth rate. We used EPCLUST (http://www.bioinf.ebc.ee/EP/EP/EPCLUST/) to perform the cluster analysis, with average linkage and a euclidian distance metric.

## Discussion

We have presented a coupled dynamic multi-species dynamic FBA and mass-transfer model of the gut microbiota. We first studied a non-spatial variant of the model, in order to determine to what extent cross-feeding can emerge in a non-evolving, diverse population of metabacteria. The individual metabacteria in this model contain the major carbohydrate fermentation pathways in the colon. Starting from glucose as a primary resource, the model produced acetate, butyrate, carbon dioxide, ethanol, formate, lactate and propionate. These fermentation products compared well with the short-chain fatty acids in the colon [35] or in an *in vitro* model of the colon [36]. Interestingly, these fermentation products were found only if the model was run with FBAwMC and not with standard FBA. This indicates that the individual metabacteria must be able to exhibit diauxic shifts, which are due to rate-yield metabolic trade-offs in FBAwMC [33, 30]. It has been argued that metabolic trade-offs in combination with mutational dynamics may already explain population diversity, even in the absence of cross-feeding or spatial heterogeneity [49]. This is an interesting finding, that may partly explain the microbial diversity in the gut. However, it is known that cross-feeding interactions exist in the gut [50, 51] and are likely to be an important factor in determining microbial diversity. Also, the results depended on cross-feeding: In the non-spatial variant of the model, only 60% of the carbon consumed by the bacteria came directly from glucose, and single metabacteria did not produce the same set of metabolites as found also *in vitro* and *in vivo*. We next ran a spatial variant of the model in a gut-like environment, a tube in which the metabolites diffuse and advect from input to output, and the bacteria attach to the gut wall. This spatially explicit, sdFBA model was extended with models of bacterial population dynamics, and ’mutation’ of the metabacteria due to the gain and loss of pathways from the local population, *e.g.*, to loss or gain of species or horizontal gene transfer. In this model, a stratified structure of metabolic niches formed, with glucose consumers in front, followed by strata inhabited by secondary and tertiary consumers.

**Figure 8.**
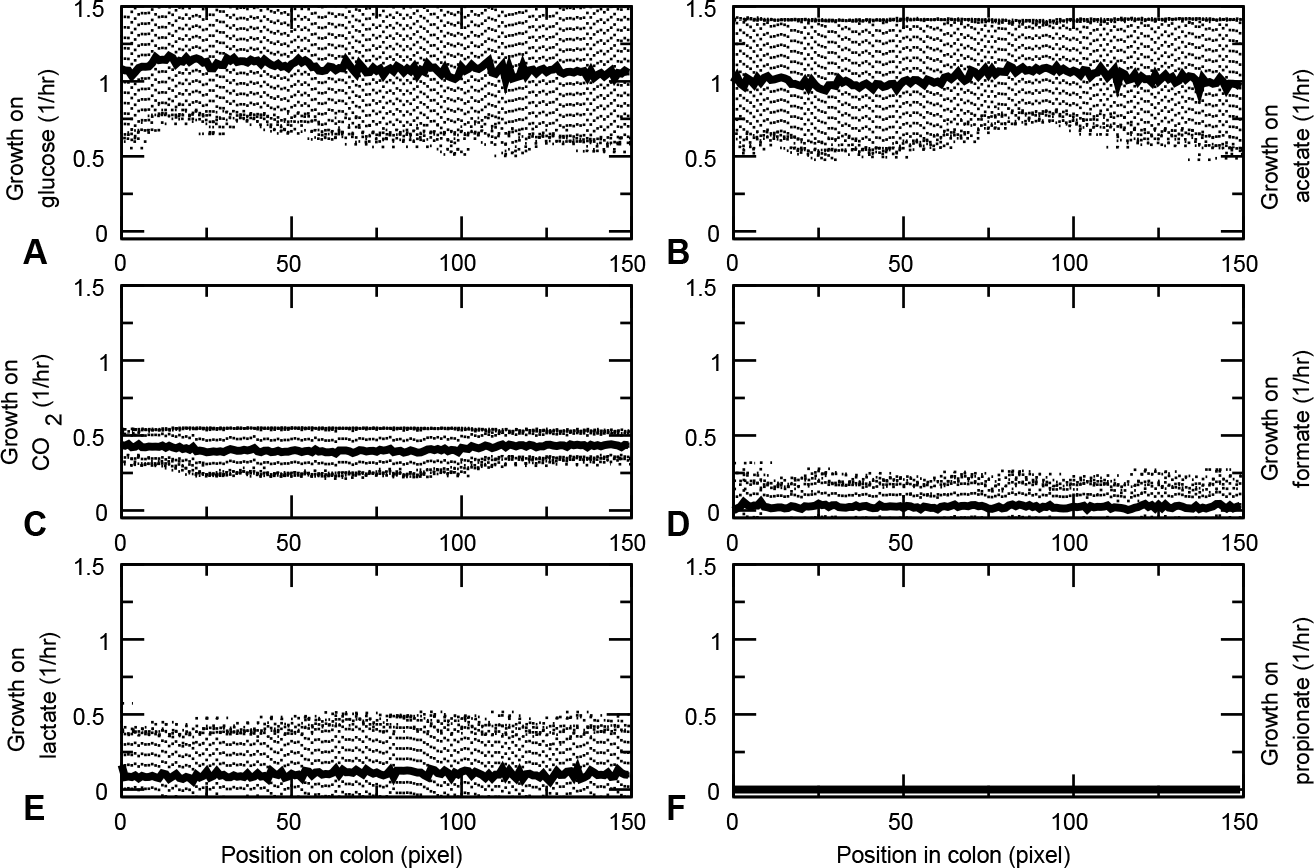
Average growth rates along the colon, when cells flowthrough the colon as fast as metabolites. Average are taken over the final part of the simulations (3500-4000 hrs) All growth rates ar calculated in the presence of unlimited hydrogen gas, water, sodium, ammonium, phosphate, sulfate and protons. (*A*) Growth rate on glucose. (*B*) Growth rate on acetate. (*C*) Growth rate on CO_2_. (*D*) Growth rate on formate. (*E*) Growth rate on lactate. (*F*)Growth rate on propionate.

Interestingly, the metabacteria specialized to their metabolic niche: Metabacteria sampled from the rear end of the tube could no longer grow on the primary resource glucose (Figure 6A), and they grew better on the secondary metabolite lactate than bacteria from the front did (Figure 6E1). This specialization was mostly due to “gene loss”, *i.e.,* simplification of the metabolic networks. Interestingly, metabacteria with reduced genomes did not have a growth advantage in our model, yet they lost essential pathways required for metabolizing the primary resource. Such “trait loss without loss of function due to provision of resources by ecological interactions” [52] is indicative of an evolutionary mechanism known as *compensated trait loss* [52]. Note, however, that because smaller metabacteria did not have a growth advantage in our model, the gene loss in our model is due to drift. Hence it differs from the Black Queen Hypothesis [53], which proposes that the saving of resources associated with gene loss accelerate the evolution of compensated trait loss. An interesting future extension of the model would consider the metabolic costs associated with the maintenance of metabolic pathways.

**Figure 9.**
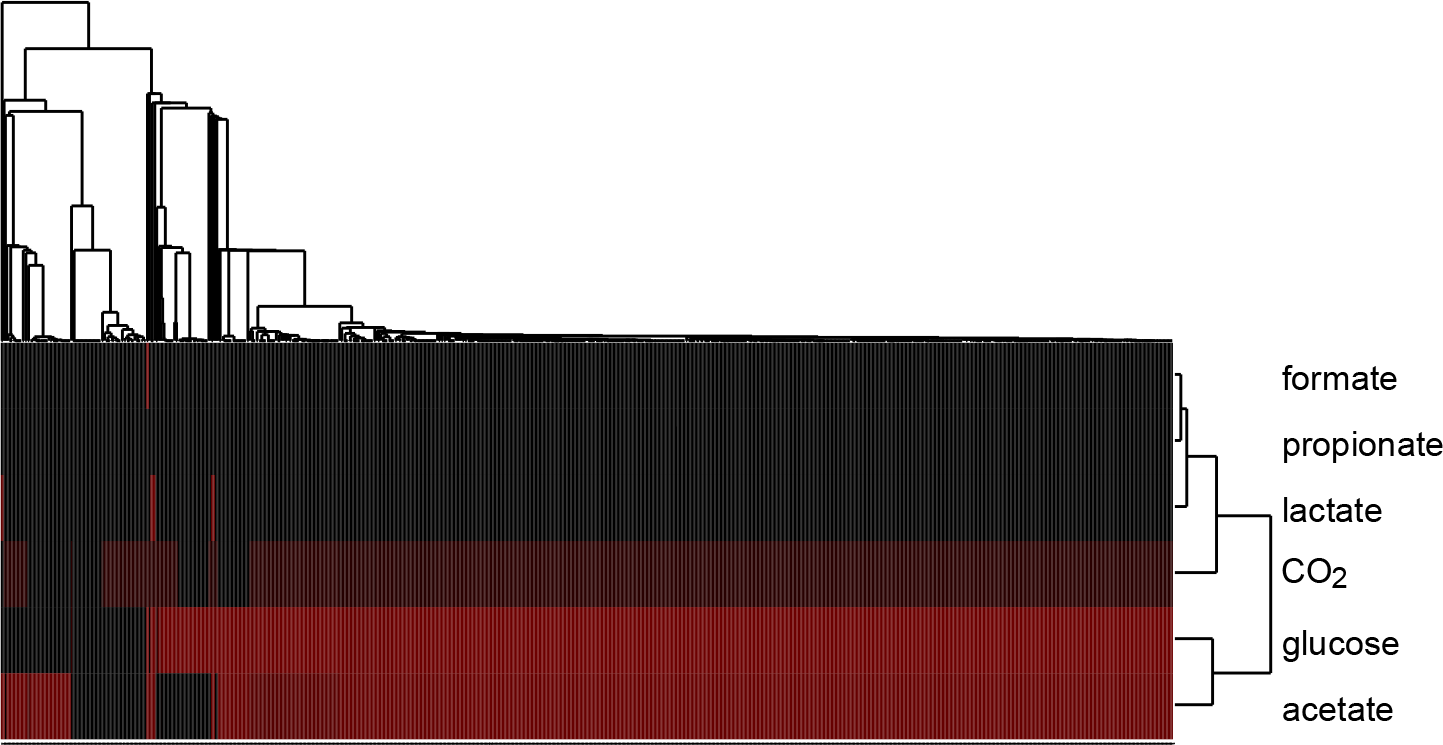
Hierarchical clustering of all cells present at the end of the simulation with cell flow, with respect to the growth rates on formate, CO_2_, propionate, lactate, glucose and acetate. Black indicates low growth rate, red high growth rate. We used EPCLUST (http://www.bioinf.ebc.ee/EP/EP/EPCLUST/) to perform the cluster analysis, with average linkage and a euclidian distance metric.

The formation of metabolic niches and the observed compensated trait loss required that the metabacteria can maintain their approximate position in the gut-like tube, *e.g.*, by adhering to the gut wall or by sufficiently fast reproduction [48]. The microbial diversification did not occur if the metabacteria moved along with the flow of the metabolites, a situation resembling diarrhea. Microbial diversity is often seen causative for diarrhea, *e.g.*, because it facilitates colonization by pathogenic species including *Clostridium difficile* [54]. Our model results suggest an additional, inverse causation, where accelerated transit reduces microbial diversity. Experimental studies are consistent with the idea that transit speed is causative for reduced diversity, but with a different mechanism: Microbiota sampled from softer stools (*i.e.*, higher BSS and faster transit time) have higher growth potential, suggesting that faster transits favor fast growing species [48]. A second potential strategy to preventing wash-out from the gut at high transit times is adherence to the gut wall *e.g.*, by the species of the P enterotype [48]. Thus these observations suggest that the reduction of microbial diversity at fast transits is due to selection for fast growing or adherent species. Our computational model suggests an alternative hypothesis, namely that increased transit times reduce the potential for bacterial cross-feeding, thus reducing the build-up of metabolic niches in the environment.

Our multiscale simulations demonstrated how cross-feeding can give rise to diverse microbial ecosystems in a gut-like geometry, but of course our model is a simplification and lacks many key features of the gut microbiota and of the gut itself. The metabacterium only contains a minimal subset of the metabolic pathways that are found in the gut microbiota. Future versions of our model could extend the current metabacterium model with additional metabolic pathways, *e.g.*, methanogenesis or sulfate reduction. Adding multiple pathways would increase the number of potential cross-feeding interactions and improve the biological realism of the model. An alternative route is to include multiple, alternative metabacteria, each representing a functional group in the human gut microbiota [55]. This would allow us to compare the metabolic diversification observed in our computational model with metagenomics data, or use the model to compare alternative enterotypes [56]. A further simplification of this first study of our model, is that we have focused exclusively on glucose metabolism. Future versions of the model will also consider lipid and amino acid metabolism, allowing us to compare the effect of alternative “diets” and consider the break-down of complex polysaccharides present in plant-derived food fibers. Future extensions of this model will also include more complex interactions with the gut wall, which is currently impenetrable as in some in *vitro* models of the gut microbiota [57, 58]. Additional terms in Eq. 6 will allow us to study the effects of SCFA from the gut lumen, and effects of the production of mucus by the gut wall [59].

## Methods

### Metabolic model

We converted the genome-scale metabolic network of *L. plantarum* [29] to a stoichiometric matrix, S. Reversible reactions were replaced by a forward and a backward irreversible reactions. Next, we added four metabolic pathways that are crucial in carbohydrate fermentation in the colon, but are not present in the network: propionate fermentation, butyrate fermentation, the acrylate pathway and the Wood-Ljungdahl pathway. We used the Kegg database (http://www.genome.jp/kegg) [60] to add the necessary reactions. For the Wood-Ljungdahl pathway, we followed the review paper [45]. Additional File 1 lists all reactions and metabolites of the genome-sscale model, in particular those that we added to the genome-scale metabolic network of *L. plantarum*.

To calculate the fluxes through the metabolic network as a function of the extracellular environment, we used flux-balance analysis with molecular crowding (FBAwMC) [33, 34]. FBAwMC assumes that all reactions through a are in steady state:

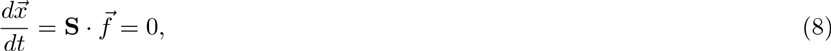

where 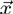 is a vector of all metabolites, 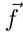 is a vector describing the metabolic flux through each reaction in the network, and S is the stoichiometric matrix. FBAwMC attempts to find a solution 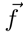 of Eq. 8 that optimizes for an objective function under a set of constraints 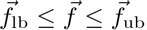 with 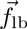 and 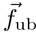 the lower and upper bounds of the fluxes. Furthermore, FBAwMC constrains the amount of metabolic enzymes in the cell. This leads to the following constraint

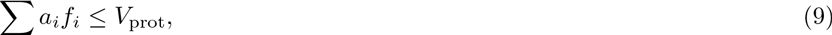

where 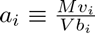 is the “crowding coefficient”, *M* the cell mass, *V* the cell volume, *v_i_* the molar volume of the enzyme catalysing reaction *i* and *b_i_* is a parameter describing the proportionality between enzyme concentration and flux. For a derivation of Eq. 9 see Ref. [33]. *V*_prot_ is a constant (0 ≤ *V*_prot_ ≤ 1) representing the volume fraction of macromolecules devoted to metabolic enzymes. We use a value of *V*_prot_ = 0.2, equal to the value used in [30] for other bacteria.

The crowding coefficients are not known for every reaction in the metabolic network. Therefore, following Vazquez and coworkers [34], crowding coefficients were chosen at random from a distribution of known crowding coefficients for *E. coli* based on published molar volumes (Metacyc [61]) and turnover numbers (Brenda [42]). Both in the well-mixed simulations as in the spatially explicit simulations, we allowed for unlimited influx of hydrogen gas, water, sodium, ammonium, phosphate, sulfate and protons. To calculate the growth rate, we find a solution of Eq. 8 that maximizes the rate of ATP production, given the crowding constraint (Eq. 9). The growth rate, *μ*, then follows from an auxiliary, empirical biomass reaction, which converts metabolic precursors into biomass (see Ref. [62] for a good introduction to FBA).

### Well-mixed model

Simulations of the well-mixed model are performed in Matlab, using the COBRA Toolbox [63]. We use an approach similar to Ref. [21] to model a population of cells in a well-mixed environment. We initiated 1000 cells with crowding coefficients for all their reactions set according to the experimental distribution of *E. coli* (see Section Metabolic model) We start with a total biomass concentration (*B*) of 0.01 gram dry weight/liter (gDW/l), divided equally over all 1000 metabacteria (*i.e.*, ∀*i* ∈ [1,1000]: *B_i_*(0) = 10^−5^ gr DW/l). At time t=0 we initiate the environment with a glucose concentration of 1.0 mM. At every time-step, the maximal uptake rate for each metabolite *j* is a function of its concentration, *c_j_* (*t*), as,

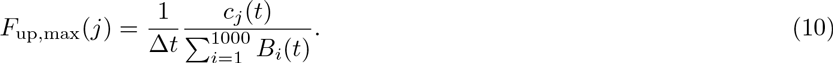

We then perform FBAwMC for all 1000 supra-bacteria and update the concentrations of all metabolites that are excreted or taken up, as,

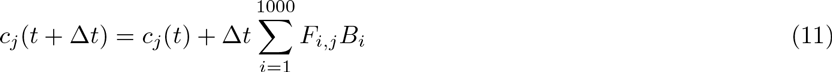

FBAwMC yields a growth rate *μ_i_* for each supra-bacterium *i*, which is used to update the biomass as,

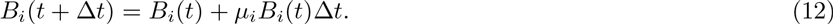

This procedure is continued until the supra-bacteria have stopped growing.

**Figure 10.**
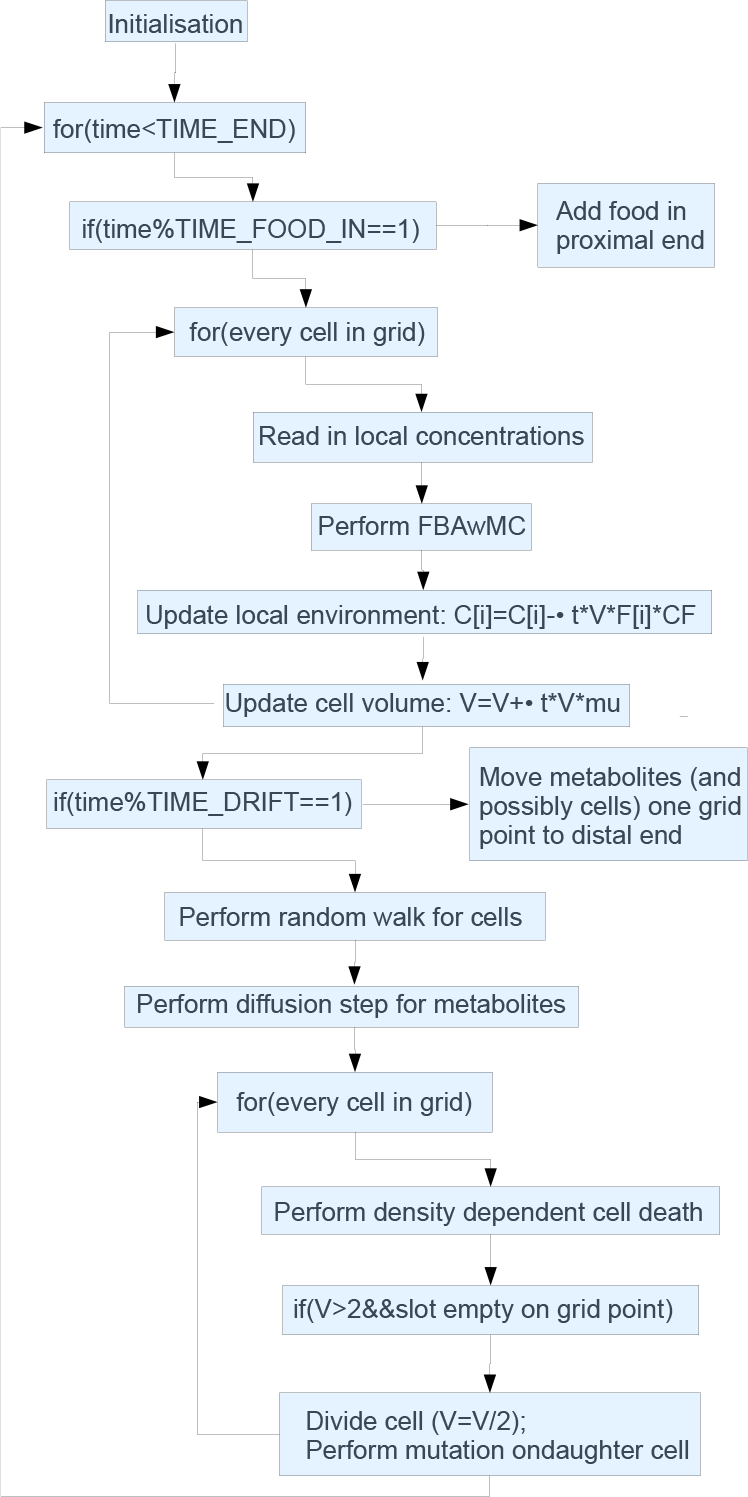
Pseudocode of the spatially explicit computational model.

### Spatially explicit, evolutionary model

For the spatially explicit simulations, we developed a C++ package to perform constraint-based modeling using the GNU Linear Programming Kit (GLPK, http://www.gnu.org/software/glpk/) as linear programming tool. The multiscale, computational model of the gut microbiota was also developed in C++. It describes individual metabacteria, or “cells” living on a grid, each with its own genome-scale metabolic network. Food enters the grid in one end, flows through the grid, diffuses over the grid and is consumed by the cells. Uptake and excretion of metabolites is calculated using the genome-scale metabolic network in each cell. The cells divide proportional to the calculated ATP production rate and mutate upon division. We simulate a total time of 4000 h (equivalent to 80000 time steps). A model description in pseudocode is given in Fig. 10. All parameters in the model are given in Table 1.

#### Initialization

We initialize the grid with cells that have the same metabolic network as in the well-mixed simulations. We choose the crowding coefficients for each reaction randomly. We allow maximally 2 cells to be present on each grid point. Thus, per grid point there are two “slots” that can be empty or filled by a cell. At time t=0, we initialize every slot of every grid point with a probability of 50% with a cell with random crowding coefficients. Because of the modeled population size (in the order of 1000 cells), each cell should be viewed as a metapopulation of bacteria that is representative for the local composition of the intestinal microbiota: i.e, a metabacterium.

**Table 1.** Parameters of the spatially explicit model.

| parameter | value | units | comments |
| --- | --- | --- | --- |
| $\Delta t$ | 3.0 | min | |
| $\Delta x$ | 1.0 | cm | |
| grid length | 150 | grid sites |  |
| grid height | 10 | grid sites |  |
| TIME_END | 4000 | hr |  |
| # slots per grid point | 2 |  |  |
| DENS_MAX | 1.0 | $\text{g DW} \cdot \text{l}^{-1}$ | see main text |
| initial density | 50% |  | assumed |
| TIME_FOOD | 8 | hr | assumed |
| FOOD_IN | 42 | mmol | assumed |
| Diffusion constant | 700 | $\mu\text{m}^2/\text{s}$ | assumed (compare $900 \mu\text{m}^2/\text{s}$ glucose in water) |
| P_MOVECELL | 0.05 |  | assumed |
| DEATH_BASAL | 0.025 | $\text{hr}^{-1}$ | assumed |
| DEATH_DENS | 2.0 | $\text{hr}^{-1}$ | assumed |
| TIME_DRIFT | 15 | min | passage time of approximately 40 hrs |
| P_CELL_FLOW | variable |  |  |
| UPTAKE_HOST | variable |  |  |
| $\mu_{\text{DEL}}$ | 0.002 | | assumed |
| $\mu_{\text{BIRTH}}$ | 0.0002 | | assumed |
| $\mu_{\text{POINT}}$ | 0.002 | | assumed |
| $\mu_{\text{POINT\_STEP}}$ | 0.2 | | assumed |

#### Food dynamics

We assumed that food enters the colon every 8 hours. In this study we consider glucose as the primary resource, because we want to focus on the bacterial diversity that can result from a single resource. Thus we assume that polysaccharides are already broken down to glucose. To allow for variability, we pick the amount of food from a normal distribution with mean of 42 mmol and a relative standard deviation of 20%. This mean value is chosen such that one the one hand all food is consumed during passage through the gut and on the other hand it allows for a sufficiently large population size (≈ 1000 metabacteria).

The glucose is consumed by the metabacteria, according to the metabolic networks. These network take into account 115 extracellular metabolites, whose dynamics are all modeled explicitly in the model. The majority of these metabolites are never produced. Production and consumption for each metabolite is modeled using

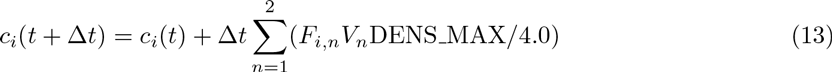

Thus, the concentration *c_i_*(*t*) of each metabolite *i* is updated each timestep Δ*t* according to the calculated influx/efflux, *F_i,n_*, and cell volume, *V_n_*, of the cells on the grid point (maximally 2). Fluxes in the metabolic network have unit mmol · g DW^−1^ · h^−1^, where external metabolite concentrations are in mmol · l^−1^.

To convert the fluxes to extracellular concentration changes, we therefore multiply with DENS_MAX; it is the maximum bacterial density in g DW · l^−1^, which is estimated as explained in Table 1. The division by four is because there can be at maximum 2 cells of volume 2 at one grid point. DENS_MAX is the maximum local density of bacterial cells; it is used to calculate the change in metabolite concentration based on the metabolite influx and efflux. If a grid point is fully occupied with two meta-bacteria the cell density at that point equals DENS_MAX. A high DENS_MAX results in large changes in extracellular concentrations due to exchange fluxes. We estimated DENS_MAX using an estimated bacterial density of 10^14^ cells/l, an estimated bacterial cell size of 10^−16^ l/cell and a cellular density of 100 g DW/l, *i.e.*, 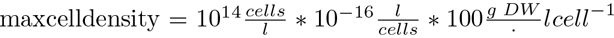[64, 65]. To prevent negative concentrations, the uptake per time step Δt is capped at

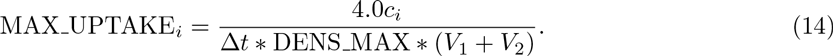

Each metabolite flows through the colon: Every 15 minutes, all metabolites are shifted one grid point to the right. This results in a passage time of 37.5 hour, similar to observed colonic transit times (*e.g.*, 39 hrs in [66]). Every metabolite also diffuses. We use a diffusion constant of 700 *μm^2^/s* for all metabolites, somewhat slower than the diffusion constant of glucose in water.

#### Population dynamics

FBAwMC yields growth rate, *μ*, for each metabacterium *i* using an empirical, auxiliary reaction [62]. The volume of the metabacterium is then updated, as

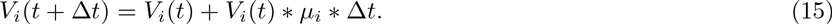

Cell death is taken into account in a density dependent way. This stabilizes the population, making sure that the population does not grow too fast if too much food is given or dies out if too little food is given. The death rate of a cell is calculated as follows

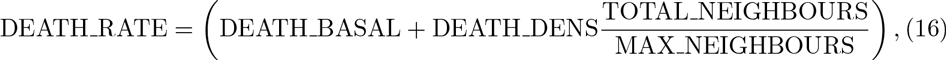

where TOTAL_NEIGHBOURS is the total amount of neighbours and MAX_NEIGHBOURS the maximum amount of neighbours (17 in the centre of the grid, because there are 2 slots per grid point).

Next the metabacteria expand into the empty patch on the same grid point when their volume exceeds a value of 2. The volume of the parent metabacterium is then equally distributed over the two daughter metabacteria. During this expansion, three types of “mutations” can occur:

a. the complete deletion of a reaction, *i.e.*, extinction of the species responsible for this reaction, with probability *μ*_DEL;
b. the reintroduction of metabolic pathways, corresponding to the invasion of the bacterium previously responsible for this pathway, with probability *μ*_BIRTH;
c. the strengthening or weakening of one of the pathways, corresponding to the relative growth or suppression of a bacterial species in the metapopulation, with probability *μ*_POINT.

To delete reaction (a) the maximal flux through that reaction is set to 0. To reintroduce a reaction (b), we release the constraint by setting it to a practically infinite value (999999 mmol/gr DW hr). A point mutation (c) corresponds to a change of the crowding coefficient (*a_i_* in Eq. 9) of that specific reaction, as

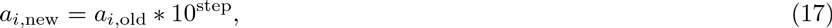

In this way, the metabacteria specialize on certain reactions, *i.e.*, by having only one or a few bacterial species in the patch. step is selected at random from a normal distribution with mean 0 and standard deviation *μ_POINT_STEP*. In this way, if the crowding coefficient is large, the mutation step will be large as well. This is necessary, because crowding coefficients are almost distributed log-normally [34, 30].

A possible non-physical side effect of this approach is that all crowding coefficients evolve to a value of *a_i_* = 0, in which case the growth rates would no longer be limited by enzymatic efficiency and volume of the patch. In reality, bacteria must trade off growth rate and growth yield (see Figure 11 and Refs.[43, 44]). To take this trade-off into account, we first calculate the total carbon uptake rate using FBAwMC as described above. We then calculate the maximal allowed growth rate, Mmax belonging to that carbon uptake rate, using the empirical formula *μ*_max_ = 1/3.9G_up_ (*i.e.*, the black curve in Fig. 11). We cap the growth rate *μ* to the maximum growth rate, μ_max_.

#### Cell movement

To model the cells’ random movement over the grid, we loop over all grid points in random order. Every grid point has two “slots” that may or may not be occupied. Each slot, whether it is occupied or not, has a probability of P_MOVECELL to exchange its position with a randomly chosen slot in a randomly chosen neighboring grid point, but this only succeeds if that slot has not already moved this turn.

An advection algorithm is introduced to model the flow of bacteria along the tube, with parameter P_CELL_FLOW determining the advection velocity relative to the metabolite flux (see Section Food dynamics). At each metabolite flow step (once every 15 minutes), with probability P_CELL_FLOW all the cells shift one grid point to the right synchronously. *I.e.*, for the default value P_CELL_FLOWν0 the cells do not flow at all, whereas for P_CELL_FLOWν1 the cells flow at the same rate as the metabolites. We performed simulations with P_CELL_FLOW ∈ {0, 0.5,1}. To mimic reentry of bacterial species from the environment, we assume periodic boundary conditions: All cells that leave the distal end of the gut, enter into the proximal end.

## Declarations

### Ethics approval and consent to participate

Not applicable

### chiladies1Consent for publication

Not applicable

**Figure 11.**
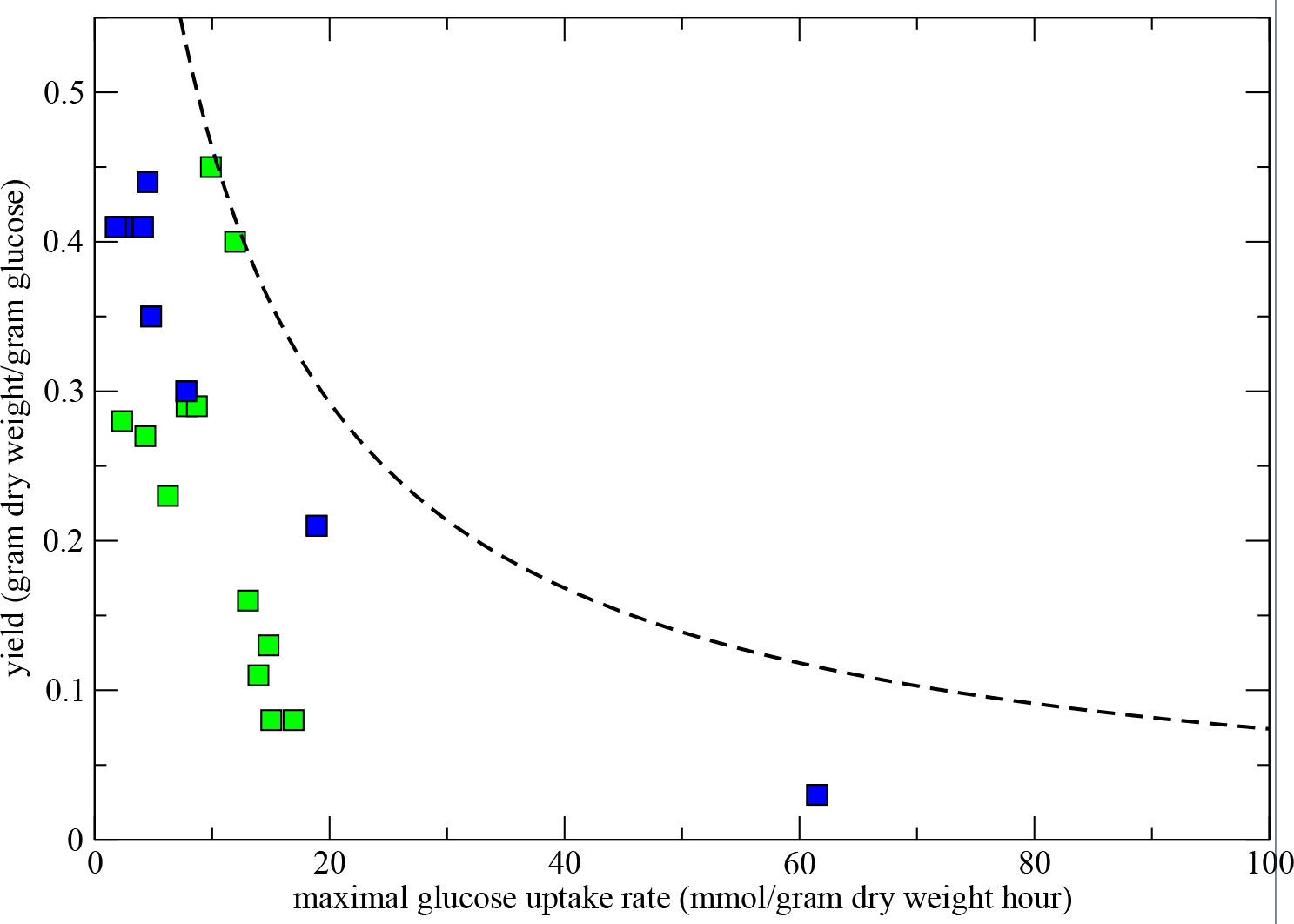
Derivation of empirical formula for maximum growth rates as a function of the glucose uptake rate. Green squares are data from yeast species [44]; blue squares represent data from bacterial species [43]. The black, dashed curve is the maximum allowed growth yield given the glucose uptake rate, *G*_up_. The empirical function is 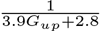 and is designed such that all data
points lie below it.

### Availability of data and material

The dataset supporting the conclusions of this article is included within the article and its additional files.”

## Author’s contributions

MvH and RM designed the study and drafted the manuscript. MvH performed the simulations and analyzed the data. All authors have read and approved the final version of the manuscript.

## Acknowledgements

We thank SURFsara (http://www.surfsara.nl) for the support in using the Lisa Compute Cluster.

## Additional Files

Additional Figure 1 — AdditionalFigure1.pdf

Simulation of the non-spatial, extended *L. plantarum* model using standard flux-balance analysis (FBA). Metabolite dynamics over time. The simulation is initialized with a pulse of glucose. Note that with standard FBA all 1000 cells behave identically, because the crowding coefficients are not used

Additional Figure 2 — AdditionalFigure2.pdf

Simulation of the non-spatial, standard *L. plantarum* model using flux-balance analysis with molecular crowding (FBA). Metabolite dynamics over time. The simulation is initialized with a pulse of glucose

Additional Figure 3 — AdditionalFigure3.pdf

Population average and standard deviation of the cross-feeding factor *C_i_* as a function of the position in the colon for all *n* = 10 runs. The averages and standard deviation are over the vertical dimension and are calculated over the final part of the simulation, from 3500 hours until 4000 hours

Additional Figure 4 — AdditionalFigure4.pdf
Population averages of the metabolite concentrations over evolutionary time of the simulation in Figure 4

Additional Figure 5 — AdditionalFigure5.pdf

Average metabolite concentraties along the tube for all *n* = 10 simulations. The averages are taken over the second half of the simulations, from 2000 hours to 4000 hours

0.1 Additional File 1 — AdditionalFile1.xls

Microsoft Excel File with all reactions and metabolites of the genome scale model of *Lactobacillus plantarum* [29], extended with proprionate fermentation, butyrate fermentation, the acrylate pathway, and the Wood-Ljungdahl pathway.

